# Standardization of postmortem human brainstem along the rostrocaudal axis to accommodate for heterogeneity in samples

**DOI:** 10.1101/2025.03.26.645559

**Authors:** Mayra Celada, Natasha Zaarour, Joni Cheung, Cassie Gross, Andrew Lim, Aron Buchman, Parastoo Saberi, Gopal Varma, Veronique VanderHorst

## Abstract

Human postmortem brain tissues provide an indispensable resource that is crucial for the understanding of neurological conditions, whether related to pathology subtype, burden, distribution or cell-type specificity. Pathology staging protocols provide guidelines for standardized sampling of brain tissues, but cover only a subset of regions affected by pathologies. Thus, to study how various neuropathologies and cell types in highly specialized circuit nodes correlate with functions specifically served by these nodes, additional protocols are necessary. This especially applies to brainstem tissues due to the small dimension of regions of interest and interindividual variability of specimens, whether due to procurement or intrinsic differences.

Here we systematically assessed factors contributing to heterogeneity in the length of whole brainstem samples and then presented a standardized approach to reproducibly assign rostrocaudal levels, with standardization relying upon readily identifiable internal landmarks. We validated this approach using postmortem MRI imaging. Standardized brainstem length correlated positively with subject height and negatively with subject age of death. By providing a reference series, reproducible levels can be assigned to individual histological sections or MRI images, i.e. when full brainstem specimens are not available and irrespective of platform, promoting reproducibility.

## Introduction

Human postmortem brain tissues provide an indispensable resource that is crucial for the understanding of neurological conditions, whether related to pathology subtypes, the degree of pathology, or to regional or cell type specific mechanisms. Sampling of brain tissues typically follows standardized protocols, in which select high yield regions are blocked and examined histologically for diagnostic and staging purposes [1–4]. While this approach serves this particular goal, it is not ideal when research questions relate to clinical-pathological studies that focus on relatively small, function-specific circuit nodes which are packed into a small volume, as is the case for the brainstem.

The brainstem connects the spinal cord with the forebrain and contains circuit nodes essential for motor control including gait, postural control, orientation and muscle tone, motor and chemosensory aspects of respiratory control, sleep and arousal, and autonomic functions including temperature, hemodynamic and bladder control, chewing and vocalization, eye-head-vestibular control, and auditory control, among others [5]. Common neuropathologies, including tauopathies, alpha-synucleinopathies, beta amyloid, TDP-43 and vascular diseases can reside in the brainstem, even in the earlier phases of neurological diseases, for review [6]. However, these pathologies are not distributed in a homogeneous way, but rather with high spatial specificity. To answer important questions about how various neuropathologies in specialized brainstem circuit nodes correlate with functions specifically served by these nodes or what drives the selective vulnerability of neuron subtypes, precise and consistent sampling of discrete regions in large cohorts is an important prerequisite for success.

Precise sampling of brainstem subregions from different decedents in larger cohorts poses several challenges. One major challenge involves inter-specimen variability in dimension of the specimens, including the rostrocaudal length. This can be due to variation in handling, blocking or sampling of tissues and/or true interindividual differences. Furthermore, (portions of) critical regions can be missed when working with standard 5mm tissue slabs, as is common in biobanking. Protocols that are developed with whole brainstems, rather than select tissue slabs, can ensure that smaller regions are not lost, and circumvent variability in sampling with tissue slabs. However, this does not solve the potential problem related to variability in specimen extraction or to intrinsic interindividual differences. Inter-specimen differences, for whichever of the above reasons, when not accounted for, inevitably result in inconsistent sampling and alignment, which can result in mismatching of regions of interest. While standard atlases of the human brain [7–11] are useful as a guide to find structures of interest, they do not help address inter-specimen differences. Experts in brainstem anatomy can navigate these complexities, but in larger cohort studies data collection and analyses are conducted by research staff with varying levels of expertise. Therefore a standardized approach is obligatory to ensure reproducible results that account for inter-specimen variability.

Here we present a method to standardize the human brainstem in terms of rostrocaudal levels using histological sections from whole brainstem specimens. We will systematically assess factors contributing to heterogeneity in whole brainstem samples across specimens and then present a standardized approach to assign reproducible levels. This approach is based upon readily identifiable internal landmarks, analogous to an approach we developed before to handle heterogeneity in lumbar spinal cord [12, 13] enabling integration of complex datasets [14]. We will validate this approach, including using postmortem MRI imaging of select whole brainstem specimens. Finally, we will explore whether brainstem length correlates with select biometric measures and demographic metrics. Through a reference series, this approach can be applied to assign reproducible levels to individual histological sections as obtained from slabs, i.e. when full brainstem specimens are not available, and to data from MRI studies.

## Materials and Methods

### Decedents

This post-mortem study includes post-mortem brainstem specimens and de-identified data from 56 decedents from the Memory and Aging Project (MAP) at Rush University in Chicago, Illinois [15]. Twenty-seven specimens were collected between 2015 and 2020 and twenty-nine from 2022 through 2024. Participants’ informed consent for tissue donation was obtained by the Rush Alzheimer’s Disease Center (RADC), and the study was approved by the Institutional Review Board (IRB) of Rush University Medical Center. The postmortem portion of the study was reviewed by the IRB at Beth Israel Deaconess Medical Center and determined to be exempt (# 2014-15/2020P000027 and #2020D001192/2020P001226).

### Brainstem specimens

Brainstem specimens were harvested through autopsy after an average postmortem interval of 10.3 hours (SD: 10.4 hrs; range 4.0-71.7hrs). A fresh sample was taken from a randomly selected side of the rostral midbrain for use in other studies and the remaining brainstem was placed in 4% paraformaldehyde. After 2 weeks, specimens were transferred to phosphate buffered saline (PBS; pH 7.4), shipped overnight to BIDMC and stored in PBS-azide at 4°C until further processing.

### Tissue processing

#### Specimen preparation

The pia mater, choroid plexus and major vessels were gently removed, and photographs of the specimens were taken from dorsal, ventral, rostral, and caudal viewpoints. We then placed a histomark along the ventral right side of the brainstem and blocked the specimens perpendicular to the floor of the 4^th^ ventricle through the mid portion of the pons, and caudal to the locus coeruleus. This was necessary to accommodate the maximum reach of the cryostat of 40mm. Similarly, if more than ∼ 0.5cm of the spinal cord was attached to the brainstem, a second cut was made caudally to the caudal end of the decussation of the corticospinal tract. The midbrain and pons from the rostral tissue block at the same side of the brainstem from which a fresh sample was removed at time of autopsy was dissected and returned to the RADC for routine neuropathology staging. We cryoprotected the remaining brainstem blocks in 25% sucrose PBS (pH 7.4) at 4°C for 2 weeks.

#### Sectioning

We used a Leica CM 1900 cryostat to collect 15 series of 50µm transverse sections in a plane perpendicular to the wall of the 4^th^ ventricle [7, 9]. The interval between consecutive sections in one series was 750µm. One case was cut at 45µm section and the same methodology was applied to this case while correcting for the difference in tissue thickness.

#### Staining

One complete series of sections from each brainstem was mounted in a gelatin solution (0.24% gelatin type A, 0.04% Chromium III potassium sulfate dodecahydrate, 0.02% sodium azide in H2O) on charged slides (12-550-15, Fisher Scientific) and air dried for at least 3 days. Slides were then washed briefly in double distilled water, dehydrated in a series of graded ethanols (5 min per solution) and subsequently defatted in xylene (2 times 30 min), 100% ethanol (2 times 20 mins), and absolute chloroform (20 mins). Slides were then rehydrated though a reverse ethanol gradient (10 min per solution), counterstained in cresyl violet (T409-25, Fisher Scientific), dehydrated in a graded series of ethanol, cleared in xylene and coverslipped using Permaslip (Alban Scientific Inc.).

#### Image processing

We ordered slides from caudal to rostral and digitized them (Hamamatsu NanoZoomer-XR; 40x). We visualized and annotated slides with NDP view 2 software 3.4. We then exported representative sections at 5x, 600 dpi to Adobe Photoshop 2024 for compilation into figures and used Adobe Illustrator 2024 to place labels. In Photoshop we adjusted orientation, contrast, brightness, and color balance of photographs of gross brainstem specimens and digitized sections and removed the area surrounding histological sections. We applied a sharpening filter to one photograph of a gross specimen.

### Postmortem MRI Image acquisition

We included postmortem MRI data from a subset of 22 specimens collected between 2022 and 2024 (P30AG072975, G.Varma) to validate the histological reconstruction approach. We imaged specimens after pia and vessel removal, but prior to blocking and cryoprotection. We placed each specimen in a PBS-filled container with a gauze to minimize movement prior to MRI in a 9.4 Tesla horizontal bore system (Bruker Biospec 94/20). A fiber optic temperature probe was inserted through the container lid to provide feedback for control of a forced air heater and maintain the specimen at a physiologically relevant target temperature of 37°C during MRI as temperature affects contrast [16]. We acquired 3D MDEFT T1-weighted and 2D multi-slice, multi-echo T2-weighted brainstem images with a slice thickness of 700µm and in-plane resolution of 350 x 350µm^2^. Images were acquired in a plane matching the plane of cryo-sectioning as much as possible. Images were processed using the scanner software (Bruker ParaVision 5.1) and analyzed using Horos v3.3.6 software. Following completion of imaging, samples were processed for histology as outlined above.

### Identification of brainstem internal landmarks and measurements

#### Gross specimens

We measured brainstem length at the side that was intact (i.e. not sampled for frozen tissue collection) of the ventral and dorsal sides of gross specimens prior to cryoprotection.

#### Calculation of absolute brainstem length from tissue sections

We calculated the length of each brainstem specimen based on the number of counterstained sections in one series of 1:15 50µm sections, with the interval between 2 consecutive sections in a series being 750µm. Only complete sections were included. As we separated the brainstem through the pons into a rostral and caudal block to accommodate the cryostat, tissue loss at the pontine level was inevitable. The degree of loss was determined by the number of incomplete sections, which accounted for a median of 2.0 sections in a 1:15 series (range 1-5; average 2.6+/- 1.0 SD), i.e. approximately 1 section for the rostral and 1 section for the caudal block. Therefore, as part of a standard approach, we added 2 sections to the mid pontine level to account for tissue loss. We put additional measures in place to adjust for any deviations from protocol, as presented in the results.

#### Identification of key landmarks in tissue sections

We used key brainstem landmarks to show the importance of standardization and to validate our standardization protocol. While the caudal end of the brainstem is classically defined by the decussation the corticospinal tract, this level was not present in all cases. The rostral end of the brainstem is classically defined by the posterior commissure, which also was not present in all cases. Therefore these intrinsic landmarks were not suitable to demarcate the rostral and caudal ends of the brainstem in this cohort study. Instead, we selected caudal and rostral landmarks that were present in all cases AND were easily identifiable even in non-counterstained sections. For the caudal landmark we chose the Obex, the point where the central canal opens into the 4^th^ ventricle. For the rostral landmark we chose the caudal end of the inferior colliculus (the most caudal section in which the inferior colliculus is well formed) in the caudal midbrain. These landmarks served to designate 0% and 100% of a relative standardization method. We chose additional landmarks for further validation of the methodology: caudal and rostral end of the decussation of the corticospinal tract (when present in the brainstem block; c-xCST and r-CST); dorsal accessory inferior olive (da-IO), caudal and rostral end of the principal inferior olive (IO); abducens nucleus (6N); pontine nulcei (PN), locus coeruleus (LC), caudal end of the decussation of the superior cerebellar peduncle (c-xSCP); rostral end of the inferior colliculus (r-IC; absent in 3 cases due to tissue handling); posterior commissure (when present in the brainstem block; PC).

#### Calculation of absolute and relative distance between landmarks in tissue sections

We determined the absolute distance between landmarks by multiplying the number of sections between landmarks (with the 2-section standard correction if landmarks were from different brainstem blocks) by the interval between 2 consecutive sections in a series (750µm). The relative level of landmarks was determined by dividing the section number in a 1:15 series relative to level of the Obex (positive if rostral, negative if caudal) by the number of sections between levels 0% and 100% (with the 2-section correction). For example, in a specimen that has a total of 53 sections between level 0% and 100%, a structure of interest that lies in the 18^th^ section rostral to the Obex was assigned a relative level of 34%.

#### Validation and optimization of leveling

A brainstem expert (NZ) made adjustments in the standard correction of 2 tissue sections based upon the position of secondary brainstem landmarks. We then assessed the impact of these additional expert corrections compared to the standard protocol. In addition, we developed criteria to detect errors in tissue handling which can impact the standardization methodology, including differences in tissue thickness or deviations in cutting angle.

#### Identification of key landmarks using MRI

We identified Obex (level 0%) and caudal end of the inferior colliculus (level 100%). While the Obex in tissue sections could be very precisely defined as the opening of the central canal, in T2-weighted MRI series, it was defined as the most caudal level at which a hypo-intense slit appeared dorsal to the central canal as the precise opening was not visible. The definition of the inferior colliculus was kept similar to the histological series. We calculated the absolute distance between key landmarks Obex and the caudal inferior colliculus based upon the absolute position of the slice from the DICOM images. The relative level for a given structure was then calculated by dividing the distance between the caudal and rostral landmarks (obex and the IC-c) by the distance between Obex and the structure of interest.

### Statistical Analyses

We compared histological and MRI derived length measurements with a two tailed t-test, after we examined normality using the D’Agostino-Pearson normality. We measured the strength of correlations between tissue and MRI data and between brainstem length and basic demographic and biometric data using the Pearson correlation coefficient. The coefficient of variability for brainstem landmarks employing the various alignment methods was calculated by dividing the standard deviation by the mean.

We assessed inter rater reliability for the critical key landmarks Obex (0%) and caudal portion of the inferior colliculus (100%), comparing the performance of an expert rater (reference) with an intermediate rater familiar with the human brainstem and two novice raters without exposure to human brainstem landmarks. We used the first 37 cases of the cohort for this validation assessment. We used GraphPad Prism 10.3.0 for all statistical analyses.

## Results

### Post-mortem human brainstem specimens vary in size

To assess the variability in brainstem length and identify the factors contributing to this, we examined 56 postmortem brainstem specimens from decedents form the Memory and Aging study. Table 1 summarizes the demographic and relevant biometric data of the decedents. Postmortem brainstem specimens varied in size (Fig. 1; Table 1). Length of gross specimen blocks ranged from 57 to 94 mm with an average of 71.4mm +/- 8.0 mm (SD) when measured from the dorsal surface and a range from 53 to 90mm with an average of 70.5 mm +/- 8.7 mm (SD) when measured from the ventral surface.

**Figure 1:**
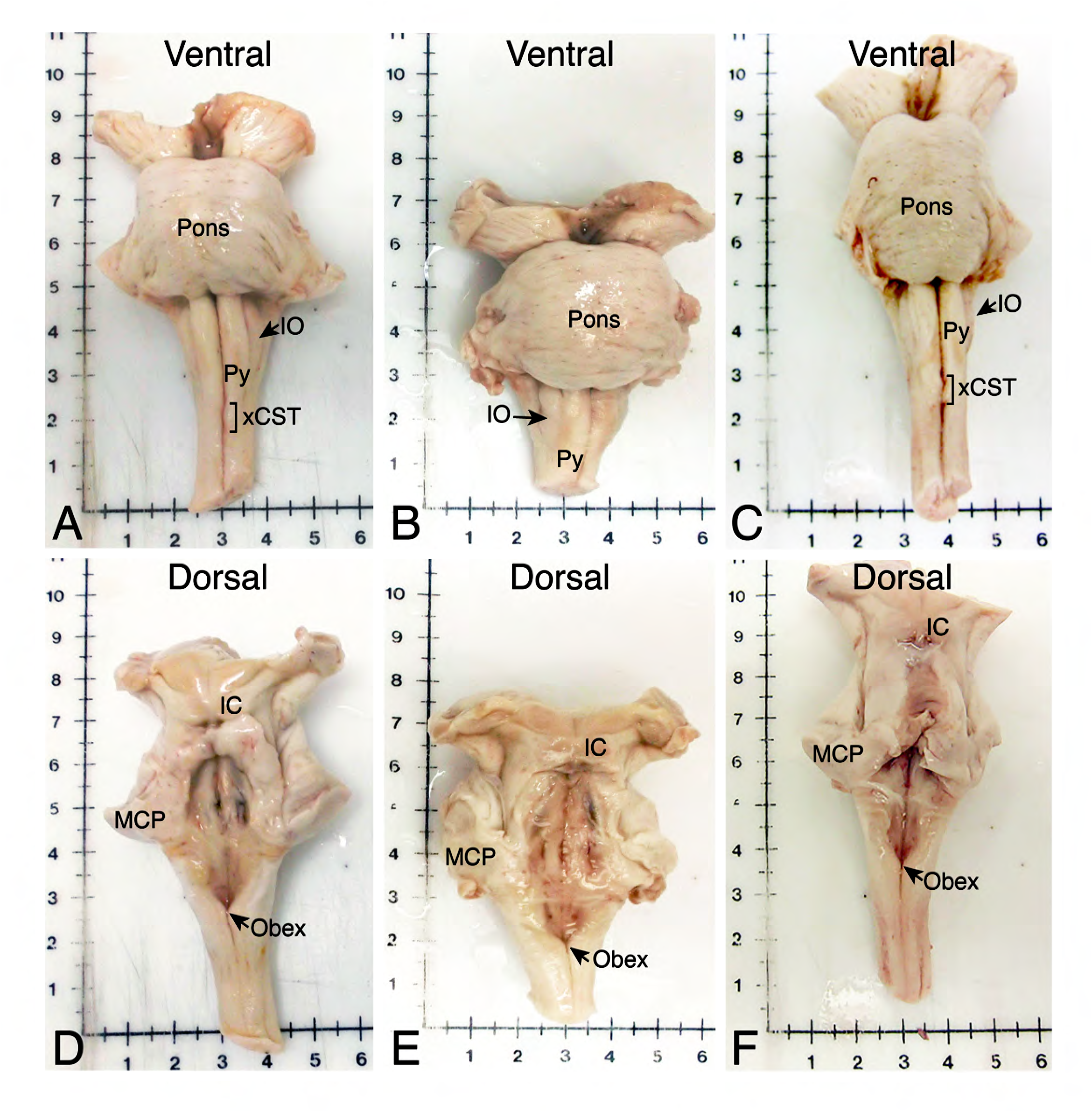
Human brainstem specimens vary in size. Photomicrographs of gross brainstem specimens. Note the variability in size due to the extraction process as well as intrinsic differences in brainstem shape. A and D: female, age at death 85 years, no cognitive impairment. B and E: female, age at death >89 years, clinical diagnosis of dementia. C and F: male, age at death > 89 years, clinical diagnosis of dementia. xCST: decussation of the corticospinal tract; Obex: point where the central canal opens into the 4^th^ ventricle; Py: pyramidal tract; IO: inferior olive; MCP: middle cerebellar peduncle; IC: inferior colliculus.

**Table 1:** Demographic, biometric and brainstem measurements of decedents.

This inter-specimen variability may be due to differences in extraction at the time of autopsy that affect the length of the medulla oblongata caudally and the midbrain rostrally, and due to inherent inter-individual differences. To examine to what extent external and intrinsic factors contribute to inter-specimen variability in brainstem length, we reconstructed each brainstem using one 1:15 counterstained series of 50µm transverse sections. The resulting 750µm steps between consecutive sections in each series were sufficient to order sections rostrocaudally (Fig. 2). In each series we then identified key landmarks at a variety of rostrocaudal levels to assess whether their alignment improved when accounting step-by-step for external and inherent factors. Key landmarks were Obex, the point where the central canal opens up into the 4^th^ ventricle at the dorsal surface of the medulla, and caudal end of the inferior colliculus at the dorsal surface of the caudal midbrain (Fig. 3; see also Figs. 1 and 2). Accessory landmarks included caudal and rostral end of the decussation of the corticospinal tract, caudal and rostral end of the inferior olive, nucleus VI, caudal end of the decussation of the superior cerebellar peduncle, rostral end of the inferior colliculus, posterior commissure, which can all be recognized in both unstained and counterstained sections (Fig. 2).

**Figure 2:**
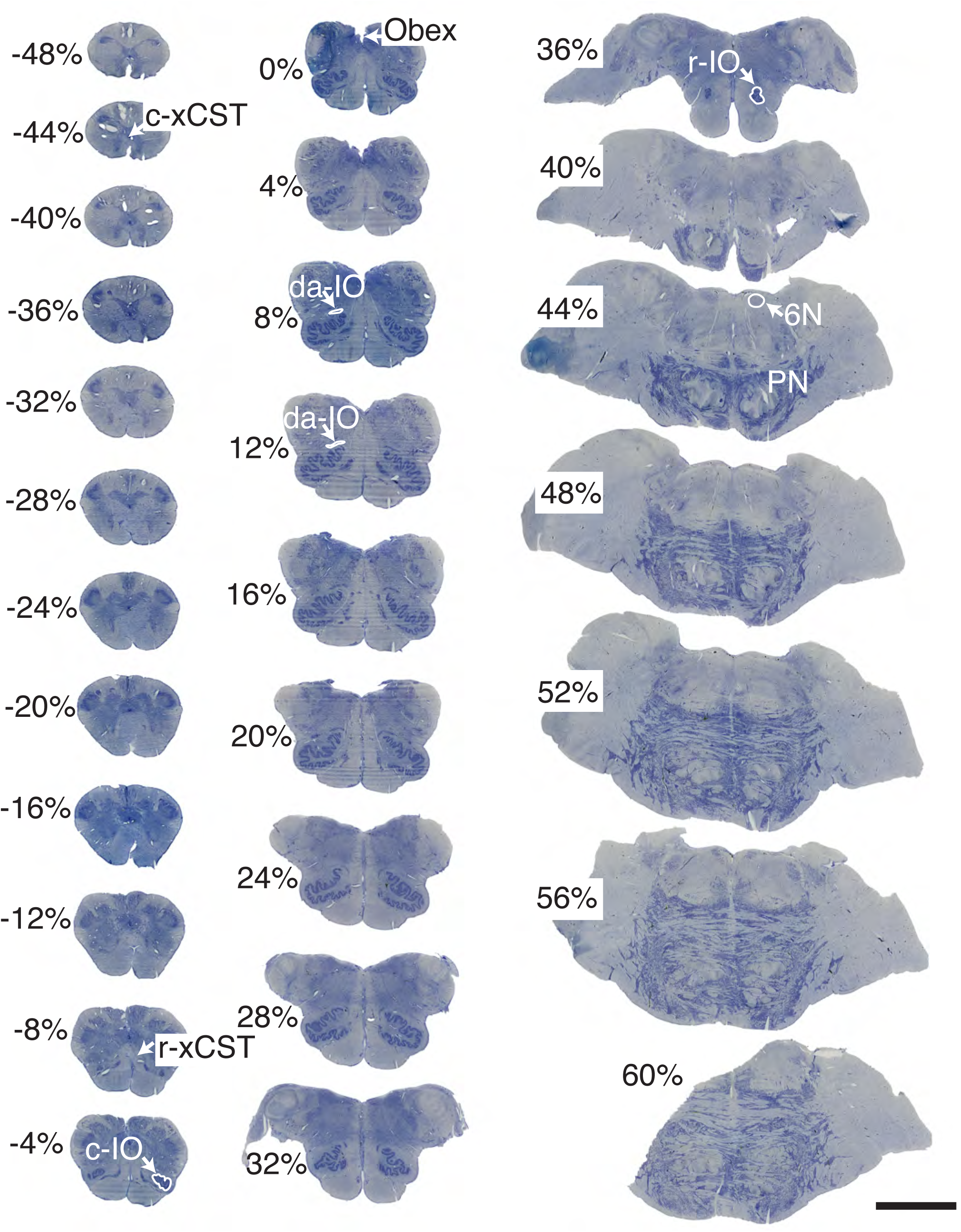
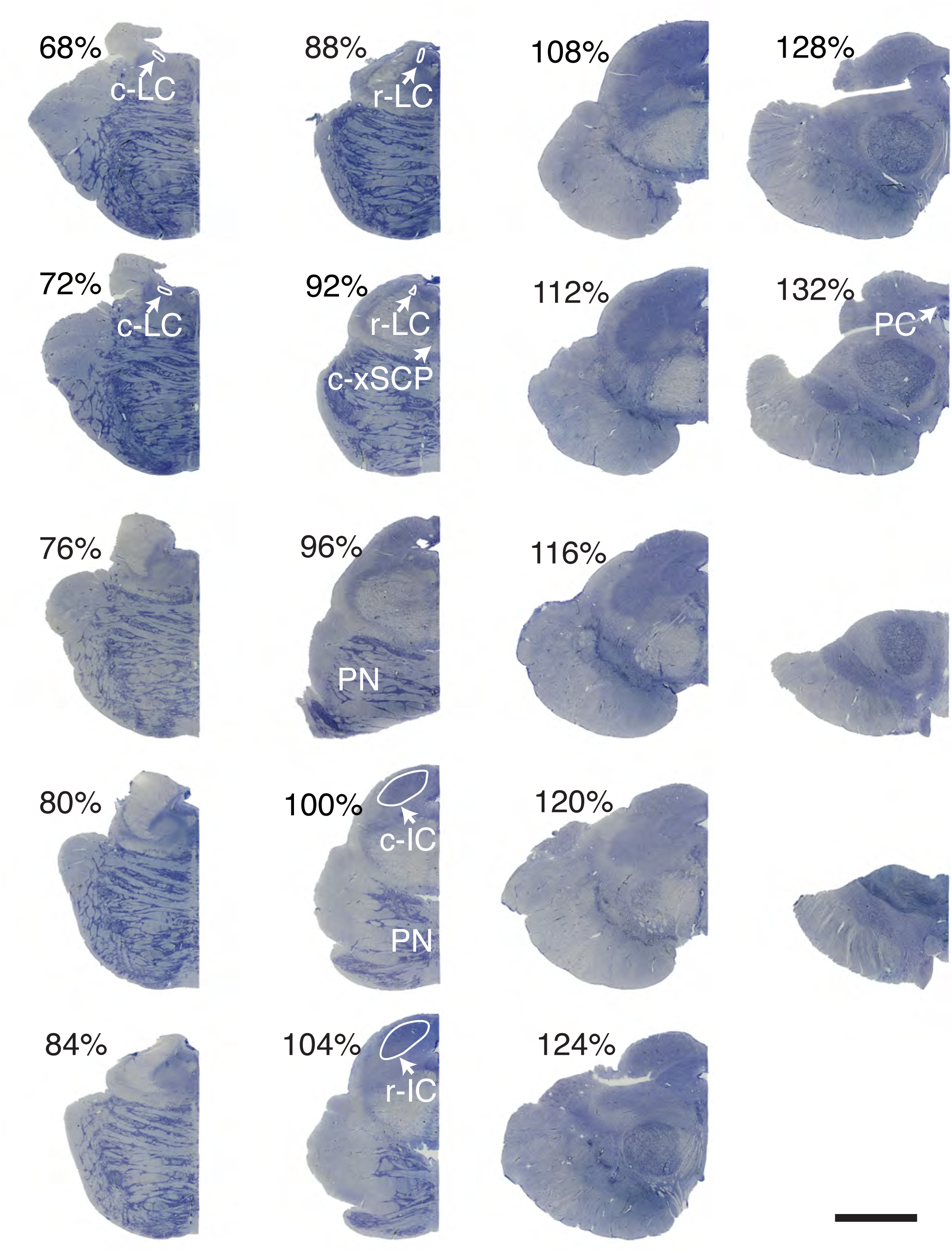
Digitized brainstem sections form a reference series. Reference series of digitized counterstained brainstem sections. Each section was cut at 50µm and each second section in a 1:15 series is represented, i.e. the interval between sections is 1.5mm. Key landmarks: inferior colliculus (IC; caudal end representing relative level 100%) and Obex (relative level 0%). Accessory landmarks: posterior commissure (PC), rostral and caudal IC (r-IC, c-IC), caudal end of the decussation of the superior cerebellar peduncle in the midline (xSCP), pontine nulcei (PN), locus coeruleus, abducens nucleus (6N), rostral and caudal end of the principle subnucleus of the inferior olive (r-IO, c-IO), dorsal accessory inferior olive (da-IO), Obex, and caudal and rostral end of the decussation of the corticospinal tract (c-xCST and r-xCST). Same case as in Fig. 1A. Please note that level 64% is not included as tissue was incomplete due to blocking of the tissue block at that level. Bar is 1 cm.

**Figure 3:**
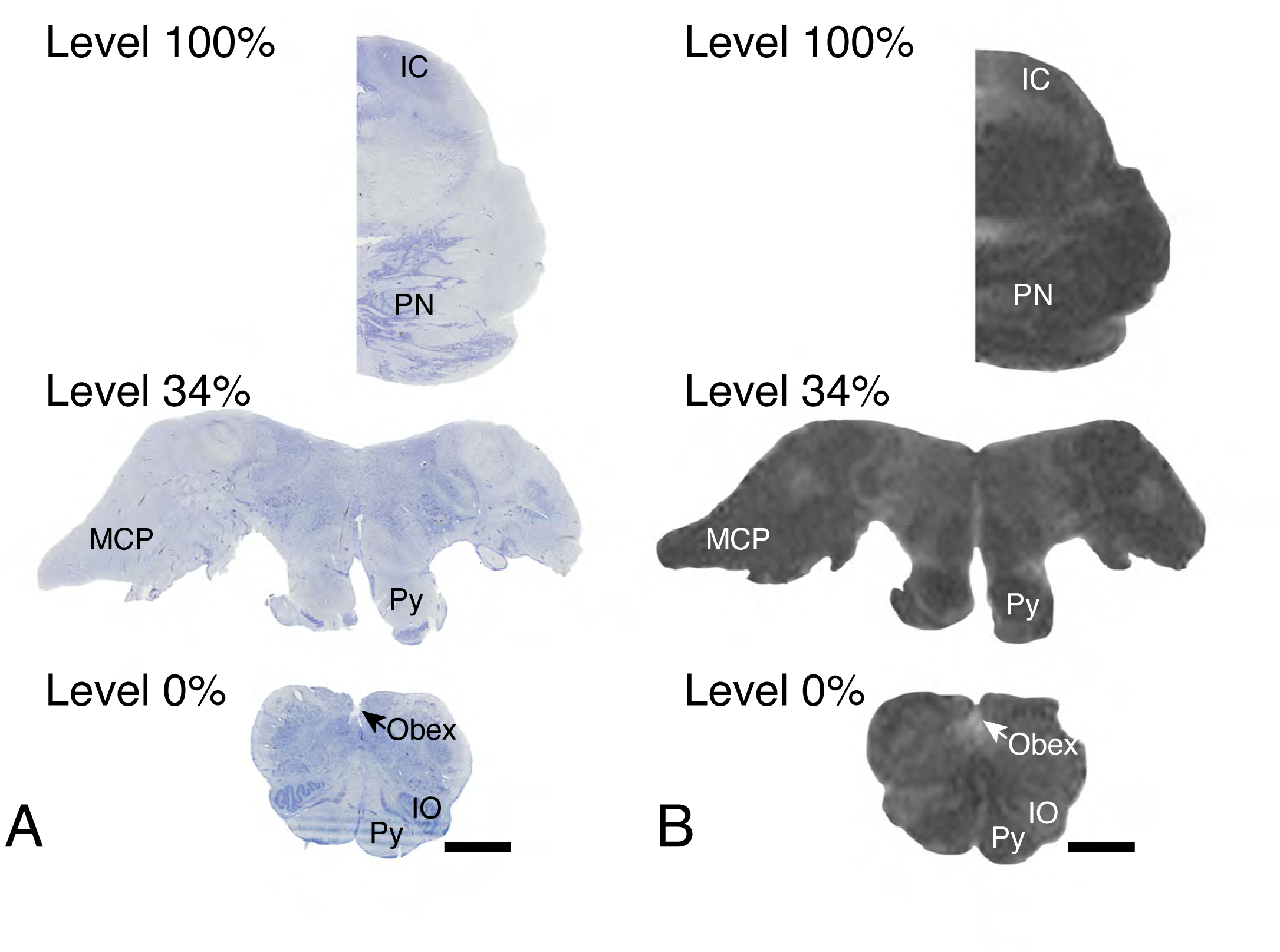
Identification of the key landmarks that designate levels 0% and 100% of a standardized relative scale in histological sections and postmortem MRI. A: Digitized sections of 50µm, transverse counterstained sections of the human brainstem (same case as in Fig. 1A) at the level of the caudal midbrain (relative level 100%, demarcated by the caudal end of the inferior colliculus, IC), the caudal pons with the rostral border of the inferior olive (in this case representing relative level 34%) and the medulla oblongata (relative level 0%, demarcated by the Obex). B: Postmortem T1-weighted images of the same brainstem as in A, representing the same levels as in A. Obex: point where the central canal opens up into the 4^th^ ventricle; Py: pyramidal tract; IO: inferior olive; MCP: middle cerebellar peduncle; PN: pontine nuclei; IC: inferior colliculus.

The length of each brainstem specimen based on the number of counterstained sections that were complete, i.e. after omitting incomplete sections at either end of the specimen, ranged from 52 mm to 80 mm (average 64.0 mm +/-6.4 mm SD; Table 1). As expected, due to the variation in harvesting, the position of landmarks was not well aligned between cases when brainstem specimens were simply aligned at their caudal edge (Fig. 4A).

**Figure 4:**
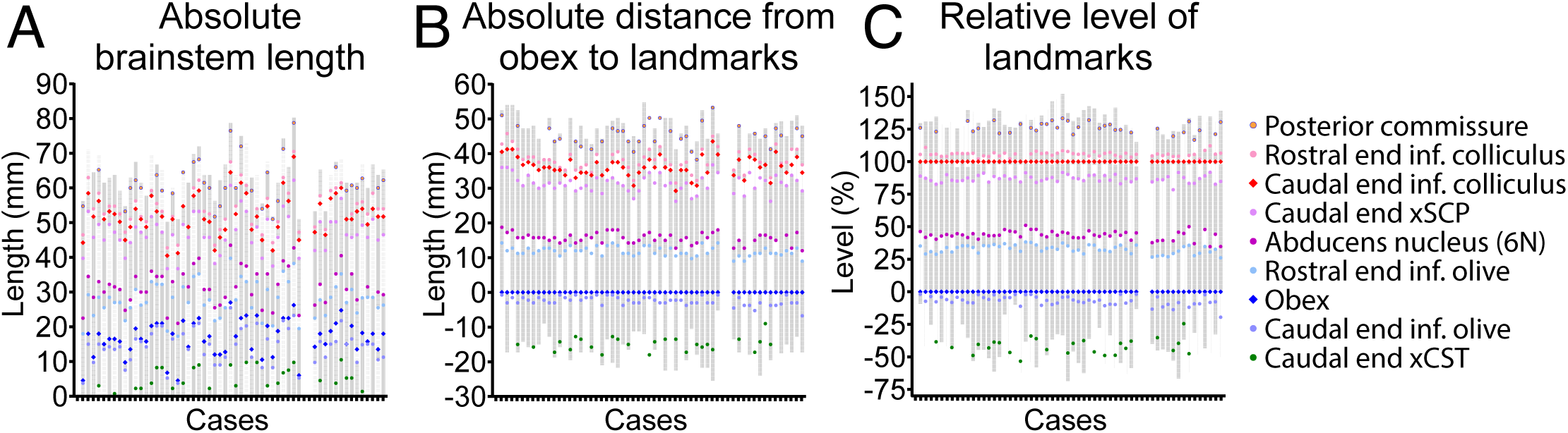
A standardized relative scale improved the inter-specimen alignment of brainstem landmarks. Length (gray area) and key landmarks (colored marks) of 56 brainstem specimens as reconstructed from 1:15 series of 50µm sections. Landmarks include the decussation of the xCST, c-IO, Obex, r-IO, xSCP, and c-IC, r-IC and PC. A: Arrangement of brainstem specimens by their caudal end resulted in poor alignment of key brainstem landmarks. B: Arrangement of brainstems according to a single caudal intrinsic landmark, the Obex, improved alignment of caudal landmarks but considerable variability remained in the rostral brainstem. C: Arrangement of the brainstems using two internal landmarks (Obex and c-IC) helped align key landmarks throughout the brainstem. Cases were grouped based upon the cutting angle (N=42 angle perpendicular to the floor of the 4^th^ ventricle; N=14 with deviation from this angle as defined in the text).

### Corrections for harvesting related factors improve but do not resolve variability in alignment between specimens

To reduce the impact of harvesting related variability, we aligned reconstructed brainstem sections based upon a single intrinsic landmark. The landmarks classically used to demarcate the caudal and rostral end of the brainstem, even though artificial, are the caudal end of the decussation of the corticospinal tract and the posterior commissure respectively. However, the classical demarcation of the caudal end of the brainstem, the decussation of the corticospinal tract was present in only 26 out of 56 cases (Figs. 1 and 4). Furthermore, the decussation itself extended rostrocaudally over 6.0 to 14.3 mm in the available specimens, making it an ambiguous landmark for the purpose of standardization. The caudal end of the posterior commissure was also only present in a subset of cases (34 out of 56; Fig. 4). These classical landmarks were therefore not suitable to serve as landmarks in cohort series with significant harvesting variability. Instead we chose the Obex as primary caudal landmark, which was present in all specimens and is readily identifiable with little training. When we aligned reconstructed brainstem series with the Obex as primary anchor (Fig. 4B), overall variability of the distribution of landmarks between cases improved, especially for landmarks close to the Obex. However, variability remained further away from this single caudal anchor, likely due to intrinsic differences in brainstem length. The wide range in absolute distance between the key caudal and rostral landmarks, i.e. Obex and inferior colliculus, from 29.3 to 43.5 mm (average 36.1mm +/- 2.9 mm SD; Table 1) supports this.

### Application of a standardized, relative scale that relies on intrinsic brainstem landmarks aligns key brainstem structures despite differences in absolute brainstem size

To manage intrinsic inter-specimen differences in brainstem size, we employed an approach similar to the one we developed to facilitate integration of datasets of the organization of motoneuron pools in the lumbar spinal cord [12, 13]. Two intrinsic landmarks, one caudal and one rostral, designate relative levels 0% and 100% of a relative scale, respectively. We used the Obex as caudal (0%) landmark and the caudal end of the inferior colliculus as the rostral landmark (100%). If the organization of structures in brainstems with different lengths is more or less fixed, the relative position of structures between landmarks is expected to be more or less stable among specimens, despite variability in absolute distance between the landmarks. Indeed this approach further improved alignment of landmarks across specimens (Fig. 4C). The coefficient of variability of positions of landmarks for the various alignment methods was highest (i.e. worst) when aligning brainstems at their caudal end and lowest (i.e. best) when applying the relative scale (Table 2).

**Table 2:** Coefficient of variability for select landmarks.

### Validation using post mortem MRI

We validated this approach using post-mortem MRI in a subset of 22 specimens. We determined the distance between the 0% and 100% key landmarks, Obex and the caudal end of the inferior colliculus, respectively (Fig. 3). Similar to the measures from histological sections, these distances varied between specimens (Fig. 5A), ranging from 30.6 to 46.5mm, with an average of 38.1 +/- 4.3 (SD) mm (Table 1). This distance, as well as the distance between other intrinsic landmarks, i.e. the rostral end of the inferior olive, correlated highly with the distances calculated from histological series (Fig. 5B). However, the distance between landmarks, both across tissue blocks (Obex to inferior colliculus) and within tissue blocks (Obex to rostral inferior olive) was smaller for the histological reconstruction compared to MRI (Fig. 5A). As this difference was even present within block, it is unlikely to be due to errors in tissue collection and most likely a consequence of the cryoprotection protocol (25% sucrose solution draws water out of specimens through osmotic pressure to avoid freeze artifacts). The relative levels assigned to accessory landmarks (i.e. rostral edge of the inferior olive and rostral edge of the inferior colliculus correlated strongly (Fig. 5C, red line) between MRI and histological series. In contrast to absolute measures, the relative position of each landmark had less variability and more closely aligned with the ideal correlation (dashed lines in Fig. 5).

**Figure 5:**
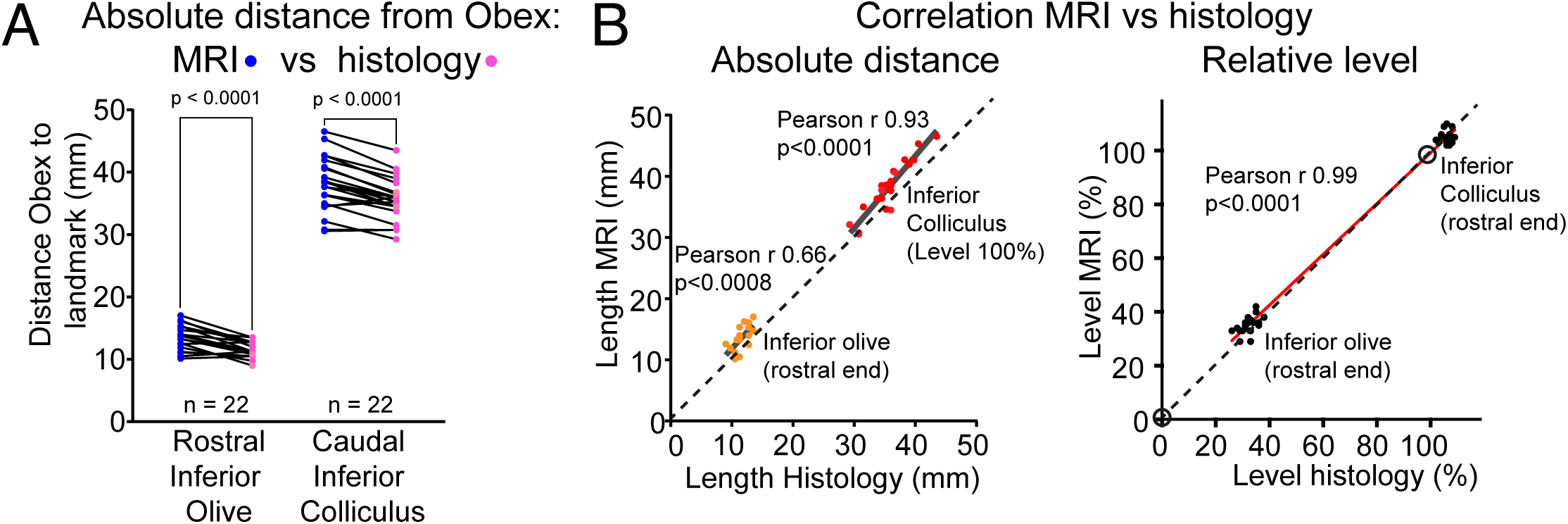
The absolute and relative position of brainstem key landmarks obtained through reconstruction of histological sections correlates well with post-mortem MRI. A: The absolute distance between landmarks was smaller in the histological series than when using MRI, i.e. T1-weighted images representing 700µm thick slices (see Fig. 3; 2-tailed paired T-test: Obex to r-IO: t = 6.47, df = 21, p < 0.0001; Obex- to c-IC: t = 6.97, df = 21, p < 0.0001). This is likely the effect of tissue shrinkage related to the cryoprotection step in the histological protocol. B: The position of key brainstem landmarks correlated well with the position derived from reconstructed sections (Pearson correlation). However, when using absolute measures, the spread within each landmark was significant due to intrinsic size differences and due to the histology shrinkage effect (see A) the regression lines shifted away from the ideal (dashed line). These errors were minimized using the relative scale for both MRI and histological sections.

### Factors that impact precision of the relative brainstem leveling approach

There are factors related to handling of brainstem specimens and cutting or imaging protocols that may interfere with applicability of the above methodology.

#### Angling of the tissue blocks

The above methodology assumes consistent sectioning or imaging in a plane perpendicular to the floor of the 4^th^ ventricle. We used landmarks in the dorsal and ventral parts of the key sections, i.e. level 0% and level 100%, to assess the impact of this variable in more detail (Figs. 2 and 3). For level 0% (Obex, a dorsal landmark), if the ventral portion represents a more rostral level than the ideal plane, the dorsal accessory inferior olive landmark will be present at the level of the Obex 0%. If the ventral portion represents a more caudal level than the ideal plane, then the principle inferior olive, ideally well developed at the level of the Obex, is absent. For level 100% (caudal end of the inferior colliculus, a dorsal landmark), if the ventral portion represents a more rostral level than the ideal plane, the substantia nigra will be well formed ventrally and the remains of the pontine nuclei are no longer present at level 100%. If the ventral portion represents a more caudal portion than the ideal plane, then the locus coeruleus may be visible and pontine nuclei are fully formed a the level of the caudal inferior colliculus. In 42 cases we found excellent angling, whereas suboptimal angles as defined above were found in 14 cases (5 being 2-4% off; 9 being >4% off).

Visualization of the relative distribution of accessory and key landmarks of cases with optimal versus deviating angles (Fig. 4, left panels versus right panels in A-C) and quantification of the coefficient of variability (Table 2) indicated that errors in cutting angle negatively affect the accuracy of the approach. If the cutting angle is poor, corrections guided by knowledge of brainstem anatomy will be necessary to align subsets of regions of interest. This should include a correction based on the expected location of a nearby accessory landmark to the anchors, e.g. if the IO is not present at level 0%, a correction is indicated to adjust c-IO to -4%, where it is expected to appear going from caudal to rostral, and this adjustment can then be applied to all ventral (but not dorsal) landmarks.

#### Correction for lost tissue

Due to blocking through the middle of the pons to accommodate the dimensions of the cryostat, tissue loss was inevitable. The number of partial sections ranged from 1-5 (average 1.9 mm +-0.8 mm SD). We employed a standard correction of 2 sections or 1.5mm to all cases (Table 1; Fig. 4A-C). We then assessed whether case by case corrections by an expert (Table 1) further improved the methodology. The impact of these changes was minimal as assessed through the coefficient of variability (Table 2). This means that unless regions of interest are very small (i.e. rostrocaudally extending for 1-3 mm only), additional adjustments are not necessary.

#### Thickness of the sections

In one case, the brainstem was cut at 45µm instead of 50µm. This did not impact the position of relative levels when we applied the correct section thickness for calculations. The standardized approach presented here can accommodate differences in thickness of tissue sections or slices and differences in the size of tissue series, as long as other parameters (i.e. sample angle) are kept controlled.

#### Raters

The relative methodological approach is reliant on correct identification of the 0% and 100% landmarks, ideally without detailed knowledge of brainstem anatomy. In the first 37 cases, we assessed the interrater reliability between an expert and 3 new raters, of which one was familiar with human brainstem sections and two had no prior exposure. All received standard instructions to identify levels 0% and 100%. Interrater reliability (Table 3) ranged from 68% to 92% for deviations of 1 or more sections (i.e. a deviation ∼1.8%) and from 97% to 100% when the error margin was set at 2 sections (i.e. a deviation of ∼3.6%). In only 1 out of 37 cases, one of the novice raters was off by 3 sections (i.e. a deviation of ∼5.4%) and none of the raters were off by more than 3 sections. From the assessment of the impact of expert corrections to account for tissue loss (see above), we already know that 1 section more or less does not impact the methodology. Altogether these findings indicate that these key structures can be easily identified among raters with minimal training.

**Table 3:** Interrater reliability for identification of key landmarks obex and inferior colliculus.

Summarizing, the use of intrinsic landmarks to standardize brainstem levels provides consistent results if basic criteria, as outlined above, are met. Being a relative scale, the approach can be adjusted for thickness and the interval between sections or slices. Through a reference series (Fig. 2 at 4% intervals; for 2% intervals see Supplemental Video), this relative scale can also be employed to assign relative levels to individual sections or slices when full brainstem series are not available, promoting reproducibility and consistency among studies.

### Does brainstem length correlate with biometric and demographic data?

We showed that the intrinsic organization of brainstem structures is more or less stable among decedents. This implicates that the distance between select caudal and rostral key landmarks (i.e. level 0% the Obex, and level 100% the inferior colliculus) can be used as a proxy for brainstem length when tissue harvesting protocols preclude measurement of the distance between classical brainstem boundaries. This enabled us to explore whether brainstem length correlated with biometric and demographic data of the decedents. Brainstem length (Obex to IC, standard correction) was larger in male than in female decedents (mean 38.0mm +/- 2.8mm SD versus 35.5mm+/-2.7 mm SD; 2 tailed t-test: t=3.08; df=54; p 0.003) and correlated with height and brain weight but not with age at death or years of education (Fig. 6). Additional clinical and pathological metrics fall outside the scope of this methodological study.

**Figure 6:**
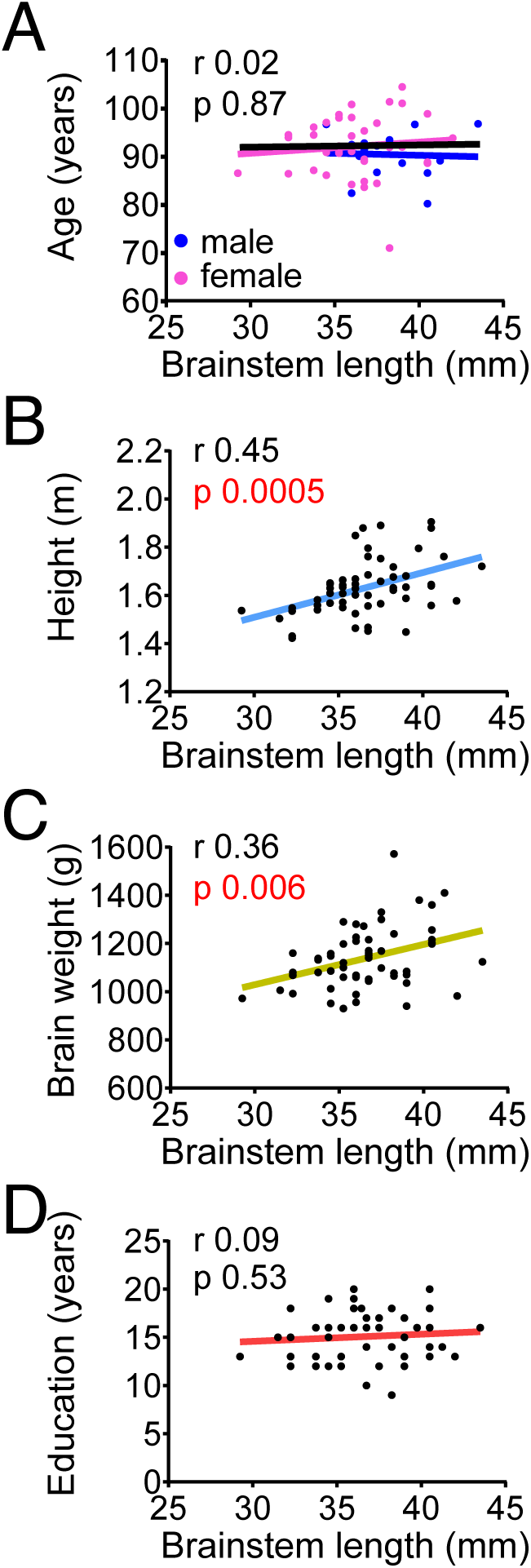
The distance between the Obex and inferior colliculus as a proxy for brainstem length correlated with body height and brain weight. Brainstem length, measured as the distance between the Obex and caudal edge of the inferior colliculus (expert correction), correlated with height (B; Pearson correlation; r 0.45; p 0.0005; N=56) and brain weight (C; Pearson correlation; r 0.36; p 0.006; N=56), but not with age at death (A; black line; Pearson correlation; r 0.02; p 0.87; N=56) or years of education (D; Pearson correlation; r 0.09; p 0.53; N=56). Correlation with clinical and pathological metrics falls beyond the scope of the present methodological study.

## Discussion

We developed a standardized approach to assign rostrocaudal levels to post mortem brainstem specimens. We developed the methodology using reconstructions from histological sections with validation using post mortem MRI from the same specimens. The approach is applicable across platforms and can be applied by staff with limited training in brainstem anatomy, if tissues are handled according to criteria that secure reproducibility. A standardized reference series of brainstem sections can be employed to assign levels when complete brainstem series are not available, as is often the case in post-mortem studies that sample tissue slabs, promoting consistency and reproducibility across studies.

### A standardized, relative leveling methodology of the human brainstem facilitates integration of data from various platforms and from larger case series

Studies that focus on human brainstem structures classically describe regions of interest based upon the anatomical position of the structure itself or of nearby landmarks or markers. While helpful, this often leaves ambiguity for several reasons that are related to both the internal organization of the brainstem and external factors. Firstly, brainstem regions of interest and landmarks are not necessarily well demarcated. While marker stains can be helpful, these are not always specific or show rather scattered stains in large fields, leaving it up to a rater, whether expert or not, to delineate the region of interest. Examples include the different components that make up the reticular formation, which extends from the caudal medulla all the way to the rostral midbrain. Secondly, brainstem structures can extend in longitudinal columns over a considerable length, especially in the medulla oblongata and pons. Examples are the midline raphe nuclei in the medulla (length ∼ 11 to 18 mm in this cohort) to and the locus coeruleus in the pons (length ∼9 to 13 mm in this cohort). While this makes it easy to find these regions, these regions are sub-organized into entities that may differ with respect to functional, connectivity, neurochemical, or molecular properties (i.e. for medullary raphe: [17–19]). Therefore, it is critical that studies refer to the sub-region that was actually sampled. Thirdly, the orientation of brainstem regions of interest can change across rostrocaudal levels, as is the case for the nucleus of the solitary tract in the medulla or locus coeruleus in the pons. As a consequence, sampling criteria may need to be adjusted depending on the level being sampled. Finally, we showed that there is significant difference in length of brainstem samples even after correction for external factors. As a result, only using absolute distance between landmarks as a guide is prone to errors.

Several external factors add to variability. This includes tissue procurement (Fig. 1). Furthermore, complete brainstem specimens are rarely available for reconstruction as tissue sections are often derived from 5mm thick routine pathology or brain bank slabs. In tissue sections derived from these slabs, smaller structures of interest are not necessarily present and for longitudinally extensive structures sampling may vary from sample to sample. This complicates feasibility or interpretation of study results when the focus is on specialized regions. A standardized methodology that is applicable across platforms, as presented in this methodological study, facilitates valid integration of data from large series of specimens, whether full brainstem samples or, via a reference series (Fig. 2), tissue slabs. This standardized system serves as the foundation for our ongoing studies, which include quantification of neuropathologies and markers in brainstem regions of interest that are not routinely sampled and for quantitative high resolution MRI.

### The principle of stability of internal landmarks

The shape and size of human and non-human primate brainstem specimens vary among individuals, in contrast to rodent models in which one can control for this variability by working with specific strains. The findings of this study demonstrated that the relative rostrocaudal location of brainstem structures are more or less preserved when controlling for differences in absolute brainstem length. This is similar to the internal organization of the spinal cord, with distinct motoneuron pools being organized functional-anatomically in a fixed relation to each other rather than based upon absolute distance or external landmarks (i.e. spinal roots; cat [12, 14], non-human and human primate[13]).

### Viewing angle is critical to consistently align internal structures

The standardized approach we presented enables one to control for differences in intrinsic absolute length between specimens. However, we showed that the cutting angle (or in MRI viewing angle) is a critical external factor that needs to be controlled for when applying the methodology to align internal structures. This is a very common but often underrecognized problem which has plagued many studies, irrespective of the species being studied, and has led to confusion or disagreement of definitions of structures. The plane applied in this study, producing the reference series, is applicable to tissues that are viewed in a plane perpendicular to the floor of the 4^th^ ventricle, as per classical studies [7, 9]. This plane is different from the plane used in the Allen Brain atlas [8] and these different planes (or any other planes) should not be used intermixed as dorsal and ventral structures will shift relative to each other between different planes. Rather, when using a different plane of sectioning, a different relative level system will be needed to align distinct regions of interest, which can be done by employing the same methodology.

In our protocol, we identified deviations from the standard plane at key levels 0 and 100, as well as control landmarks. In these flagged cases, adjustments done by an expert, including separately leveling the dorsal and the ventral part of the sections, will be necessary to extract data that aligns with the reference plane. Additional issues related to tissue handling such as excessive tissue loss at the ends of the tissue blocks should also be flagged and handled by expert staff.

### Correlation of brainstem length with demographic and biometric data

Using the absolute distance between the Obex and the caudal border of the inferior colliculus as a proxy for brainstem length, we found that brainstem length was positively correlated with height and brain weight. In this small sample of older adults, it did not correlate with age at death or education. Work is underway to explore these correlations in a larger cohort and between surface area, aging and neurodegeneration, similar to MRI studies which showed an effect of aging on regional brainstem size [20–22].

### Strengths and limitations of the study

The approach developed in this study has several strengths. It can be implemented by staff with limited knowledge of brainstem anatomy, and the reference series for relative levels enables the assignment of relative, standardized levels when complete tissue series are unavailable. The approach was further verified by excellent interrater variability, especially after proper training. Standardization will facilitate integration of large datasets or when different platforms or modalities are being integrated (fresh frozen, paraffin embedded formalin fixed, cryo formalin fixed or MRI). This methodology can be adapted to different viewing or cutting angles. Limitations of this study are that the protocol was developed with brainstems from older adults, though the same principle is expected to hold in younger populations. As is true for any methodology, errors related to tissue handling, including sectioning at an angle that deviates from protocol, sectioning at uneven thickness, and excessive tissue loss at the edges of tissue blocks all introduce additional variability. The impact of such errors can be minimized by implementation of criteria to check for deviations and guidance in adjustments. However, in case of major deviations from protocol, expert input will be necessary. Another limitation is that we only assessed the rostrocaudal dimension. While we correlated brainstem size with biometric and demographic data, this was based upon cross sectional data only and therefore the results do not inform on correlations between longitudinal biometric and brainstem data. In addition, further work is needed to map brainstem sub structures within this framework and to integrate data from larger series of specimens, including longitudinal data and multimodal structural and pathological data from histological and MRI datasets.

## Acknowledgements

The authors would like to thank Asma Hassani, Paulina Aragay Herrera-Blanc, Maja Johnson and Jane Fergusson for their assistance.

## Author contributions

-design and concept: VV, JC, CG

-first draft: VV

-analyses: JC, CG, NZ, MC, PS, GV, VV

-figures: MC, NZ, GV, PS, VV

-edit final version: JC, CG, MC, NZ, PS, AB, AL, GV, VV

## Grants

R01AG047976 (Buchman, VanderHorst);

R01AG071638 (VanderHorst; Buchman; Lim);

P30AG072975 (G.Varma)

## References

1. Braak H, Muller CM, Rub U, Ackermann H, Bratzke H, de Vos RA, et al. Pathology associated with sporadic Parkinson’s disease--where does it end? J Neural Transm Suppl. 2006;(70):89–97. doi: 10.1007/978-3-211-45295-0_15. PubMed PMID: 17017514.

2. Braak H, Alafuzoff I, Arzberger T, Kretzschmar H, Del Tredici K. Staging of Alzheimer disease-associated neurofibrillary pathology using paraffin sections and immunocytochemistry. Acta Neuropathol. 2006;112(4):389–404. Epub 20060812. doi: 10.1007/s00401-006-0127-z. PubMed PMID: 16906426; PubMed Central PMCID: PMCPMC3906709.

3. Thal DR, Rub U, Orantes M, Braak H. Phases of A beta-deposition in the human brain and its relevance for the development of AD. Neurology. 2002;58(12):1791–800. doi: 10.1212/wnl.58.12.1791. PubMed PMID: 12084879.

4. Josephs KA, Murray ME, Whitwell JL, Parisi JE, Petrucelli L, Jack CR, et al. Staging TDP-43 pathology in Alzheimer’s disease. Acta Neuropathol. 2014;127(3):441–50. Epub 20131116. doi: 10.1007/s00401-013-1211-9. PubMed PMID: 24240737; PubMed Central PMCID: PMCPMC3944799.

5. Paxinos G, Xu-Feng H, Sengul G, Watson C. Organization of Brainstem Nuclei. The Human Nervous System: Elsevier; 2012. p. 260–327.

6. Goedert M. NEURODEGENERATION. Alzheimer’s and Parkinson’s diseases: The prion concept in relation to assembled Abeta, tau, and alpha-synuclein. Science. 2015;349(6248):1255555. doi: 10.1126/science.1255555. PubMed PMID: 26250687.

7. Olszewski J, Baxter D. Cytoarchitecture of the human brain stem: Basle & New York: S. Karger; 1954 1954.

8. Shen EH, Overly CC, Jones AR. The Allen Human Brain Atlas: comprehensive gene expression mapping of the human brain. Trends in Neurosciences. 2012;35(12):711–4. doi: 10.1016/j.tins.2012.09.005.

9. Mai JK MM, Paxinos G. . Atlas of the Human Brain 4th edition ed. Amsterdam: Academic Press 2015.

10. Coulombe V, Saikali S, Goetz L, Takech MA, Philippe É, Parent A, et al. A Topographic Atlas of the Human Brainstem in the Ponto-Mesencephalic Junction Plane. Frontiers in Neuroanatomy. 2021;15:627656. doi: 10.3389/fnana.2021.627656.

11. Conti A, Gambadauro NM, Mantovani P, Picciano CP, Rosetti V, Magnani M, et al. A Brief History of Stereotactic Atlases: Their Evolution and Importance in Stereotactic Neurosurgery. Brain Sciences. 2023;13(5):830. doi: 10.3390/brainsci13050830.

12. Vanderhorst VGJM, Holstege G. Organization of lumbosacral motoneuronal cell groups innervating hindlimb, pelvic floor, and axial muscles in the cat. The Journal of Comparative Neurology. 1997;382(1):46–76. doi: 10.1002/(SICI)1096-9861(19970526)382:1<46::AID-CNE4>3.0.CO;2-K.

13. Gross C, Ellison B, Buchman AS, Terasawa E, VanderHorst VG. A novel approach for assigning levels to monkey and human lumbosacral spinal cord based on ventral horn morphology. PloS One. 2017;12(5):e0177243. doi: 10.1371/journal.pone.0177243.

14. Yakovenko S, Mushahwar V, VanderHorst V, Holstege G, Prochazka A. Spatiotemporal activation of lumbosacral motoneurons in the locomotor step cycle. J Neurophysiol. 2002;87(3):1542–53. doi: 10.1152/jn.00479.2001. PubMed PMID: 11877525.

15. Bennett DA, Buchman AS, Boyle PA, Barnes LL, Wilson RS, Schneider JA. Religious Orders Study and Rush Memory and Aging Project. Journal of Alzheimer’s Disease. 2018;64(s1):S161–S89. doi: 10.3233/JAD-179939.

16. Birkl C, Langkammer C, Krenn H, Goessler W, Ernst C, Haybaeck J, et al. Iron mapping using the temperature dependency of the magnetic susceptibility. Magn Reson Med. 2015;73(3):1282–8. Epub 20140417. doi: 10.1002/mrm.25236. PubMed PMID: 24752873.

17. Brust RD, Corcoran AE, Richerson GB, Nattie E, Dymecki SM. Functional and developmental identification of a molecular subtype of brain serotonergic neuron specialized to regulate breathing dynamics. Cell Rep. 2014;9(6):2152–65. Epub 20141211. doi: 10.1016/j.celrep.2014.11.027. PubMed PMID: 25497093; PubMed Central PMCID: PMCPMC4351711.

18. Hennessy ML, Corcoran AE, Brust RD, Chang Y, Nattie EE, Dymecki SM. Activity of Tachykinin1-Expressing Pet1 Raphe Neurons Modulates the Respiratory Chemoreflex. J Neurosci. 2017;37(7):1807–19. Epub 20170116. doi: 10.1523/JNEUROSCI.2316-16.2016. PubMed PMID: 28073937; PubMed Central PMCID: PMCPMC5320611.

19. Okaty BW, Freret ME, Rood BD, Brust RD, Hennessy ML, deBairos D, et al. Multi-Scale Molecular Deconstruction of the Serotonin Neuron System. Neuron. 2015;88(4):774–91. Epub 20151105. doi: 10.1016/j.neuron.2015.10.007. PubMed PMID: 26549332; PubMed Central PMCID: PMCPMC4809055.

20. Iglesias JE, Van Leemput K, Bhatt P, Casillas C, Dutt S, Schuff N, et al. Bayesian segmentation of brainstem structures in MRI. Neuroimage. 2015;113:184–95. Epub 20150314. doi: 10.1016/j.neuroimage.2015.02.065. PubMed PMID: 25776214; PubMed Central PMCID: PMCPMC4434226.

21. Lambert C, Chowdhury R, Fitzgerald TH, Fleming SM, Lutti A, Hutton C, et al. Characterizing aging in the human brainstem using quantitative multimodal MRI analysis. Front Hum Neurosci. 2013;7:462. Epub 20130820. doi: 10.3389/fnhum.2013.00462. PubMed PMID: 23970860; PubMed Central PMCID: PMCPMC3747448.

22. Luft AR, Skalej M, Schulz JB, Welte D, Kolb R, Burk K, et al. Patterns of age-related shrinkage in cerebellum and brainstem observed in vivo using three-dimensional MRI volumetry. Cereb Cortex. 1999;9(7):712–21. doi: 10.1093/cercor/9.7.712. PubMed PMID: 10554994.

